# Chromosome segregation fidelity is controlled by small changes in phospho-occupancy at the kinetochore-microtubule interface

**DOI:** 10.1101/2021.02.16.431549

**Authors:** Thomas J. Kucharski, Rufus Hards, Kristina M. Godek, Scott A. Gerber, Duane A. Compton

## Abstract

Kinetochore protein phosphorylation promotes the correction of erroneous microtubule attachments to ensure faithful chromosome segregation during cell division. Determining how phosphorylation executes error correction requires an understanding of whether kinetochore substrates are completely (i.e. all-or-none) or only fractionally phosphorylated. Using quantitative mass spectrometry (MS), we measured phospho-occupancy on the conserved kinetochore protein Hec1 (NDC80) that directly binds microtubules. None of the positions measured exceeded ∼50% phospho-occupancy, and the cumulative phospho-occupancy changed by only ∼20% in response to changes in microtubule attachment status. The narrow dynamic range of phospho-occupancy is maintained by ongoing phosphatase activity. Further, both Cdk1-Cyclin B1 and Aurora kinases phosphorylate Hec1 to enhance error correction in response to different types of microtubule attachment errors. Thus, networks of kinases and phosphatases maintain low inherent phospho-occupancy to promote microtubule attachment to kinetochores while providing for high sensitivity of kinetochore-microtubule attachments to very small changes in phospho-occupancy to ensure high mitotic fidelity.

## Introduction

The kinetochore is a complex protein structure that provides the link between chromosomes and microtubules during mitosis ^1^. Attachments between kinetochores and microtubules are highly regulated in order to correct attachment errors that frequently occur in early mitosis to prevent aneuploidy ^2–6^. The Knl1-Mis12-NDC80 (KMN) complex has emerged as a critical target for regulation at the kinetochore as this complex plays an essential role in microtubule attachment ^7^. The Aurora A and B kinases have been shown to promote error correction through phosphorylation of these proteins, while the PP1 and PP2A-B56 phosphatases have been shown to oppose these changes. Phosphorylation of the KMN network by these kinases reduces their affinity for microtubules, enabling kinetochore-microtubule (k-MT) detachment to promote error correction ^5, 7–11^. Therefore, it has been suggested that this network “tunes” the phosphorylation state on kinetochores to precisely regulate the stability of k-MT attachments ^3, 12^. However, there is currently no systems level understanding of how these kinases and phosphatases coordinate to phosphorylate kinetochore substrates to provide this tuning of k-MT stability. Current models suggest that phosphorylation acts as an all-or-none mechanism and it remains unknown how the concerted actions of kinases and phosphatases generate the net (sum of kinase and phosphatase activities) phosphorylation on specific sites in response to k-MT attachment errors ^4, 13^. Although plausible, it seems improbable that kinetochore phosphorylation operates through an all-or-none mechanism because saturation of phosphorylation sites on kinetochore substrates would be anticipated to severely impair k-MT attachment, and conversely, complete lack of phosphate would be anticipated to create hyper-stable k-MT attachments that impede error correction.

Hec1 has an n-terminal tail comprising the first 70 amino acids that provides a primary microtubule attachment site in kinetochores. It has been demonstrated that up to 9 sites among this short sequence are phosphorylated in human cells, yet mutational analysis has revealed that wild-type function can be provided by Hec1 possessing only 1-2 phosphorylation mimicking sites ^5–7, 14–16^. Therefore, it is unclear how to reconcile the functional sufficiency provided by only two phosphorylation sites with the evidence that nine sites serve as phosphoacceptors, and this underscores our lack of understanding of the actual phosphorylation occupancy of these sites during mitosis. Conceptually, the determination of post-translational modification site occupancy is necessary to explain the mechanisms through which these modifications regulate biological processes and, with respect to phosphorylation, to determine how kinase and phosphatase networks create the net phospho-occupancy on specific sites to “tune” phosphorylation to affect the stability of k-MT attachments. Despite the potential for mechanistic insight, occupancy analysis is not yet routine and instead, differences in phosphorylation are routinely reported as relative fold-change ^17^. However, a 2-fold change in relative phosphorylation could arise from a change in absolute phospho-occupancy of either 1% to 2% or 50% to 100%, and these two scenarios have different implications for how sensitive the system is for regulating the stability of k-MT attachments. This issue cannot be resolved using site-specific amino acid substitution because mutant proteins inherently adopt one extreme (alanine mimics zero phosphorylation occupancy) or the other (aspartic acid or glutamic acid mimic 100% phosphorylation occupancy) relative to mimicking phosphorylation status.

To investigate the phosphorylation-dependent mechanisms that govern mitotic fidelity, we selected the protein Hec1 as a candidate to measure phospho-occupancy because it is crucial for microtubule binding at the k-MT interface and because previous models posit that phosphorylation at multiple sites facilitates k-MT detachment to correct attachment errors ^5, 6, 14, 16^. We determined that Hec1 is only partially phosphorylated, due to constant phosphatase activity, at all sites queried under a variety of microtubule attachment conditions, and that only small changes in phosphorylation occupancy are associated with changes in k-MT attachment status including erroneous attachments. We also discovered that Cdk1-Cyclin B1 plays a direct role in k-MT error correction through phosphorylation of the Hec1 tail that is critical for accurate chromosome segregation. Thus, the system of kinases and phosphatases operates to limit the extent of phosphorylation to both allow for initial k-MT attachments and ensure efficient error correction.

## Results

### Hec1 is phosphorylated at only partial occupancy during mitosis

To efficiently purify Hec1, we created human cells expressing Hec1 fused to a 3XFLAG tag using CRISPR/cas9 editing to insert the tag into the endogenous locus of Hec1 (Figure S1A). Addition of the tag did not perturb Hec1 function or mitotic progression as judged by Hec1 kinetochore localization, the lagging chromosome rate in anaphase, mitotic duration or binding to the known Knl1-Mis12-NDC80 (KNL) complex member and Hec1 binding partner Nuf2 (Figure S1B-F).

We purified Hec1-3XFLAG via immunoprecipitation (Figure S2C) and analyzed it by mass spectrometry (MS) to determine phosphorylation site occupancy. Since it cannot be assumed that a phospho-peptide and cognate non-phospho-peptide will be detected equally, we monitored the change in abundance of non-phosphopeptide cognates with and without calf intestinal phosphatase (CIP) treatment to dephosphorylate half of each sample by differential, isotopically-encoded reductive dimethyl labeling. MS analysis confirmed that the CIP treatment completely dephosphorylates peptides under these conditions (Figure S2F). The untreated and CIP-treated fractions are then remixed and analyzed by MS (Figure 1A, S2D). Notably, this approach cannot distinguish site-specific occupancies for peptides containing more than one phosphorylated residue, and instead integrates their non-overlapping occupancies as a single value. To explore if phosphorylation occupancy changes under different conditions of microtubule attachment, we purified total Hec1-3XFLAG from mitotic cells lacking microtubules (500 ng/ml nocodazole (NOC)), with mitotic spindle defects that create k-MT attachment errors (100 ng/ml NOC or 25 μM *S*-Trityl-L-cysteine (STLC)) ^18^, or with predominantly correct bi-oriented k-MT attachments (25 μM proTAME) ^19^ (Figure S2A,S2B).

**Figure 1:**
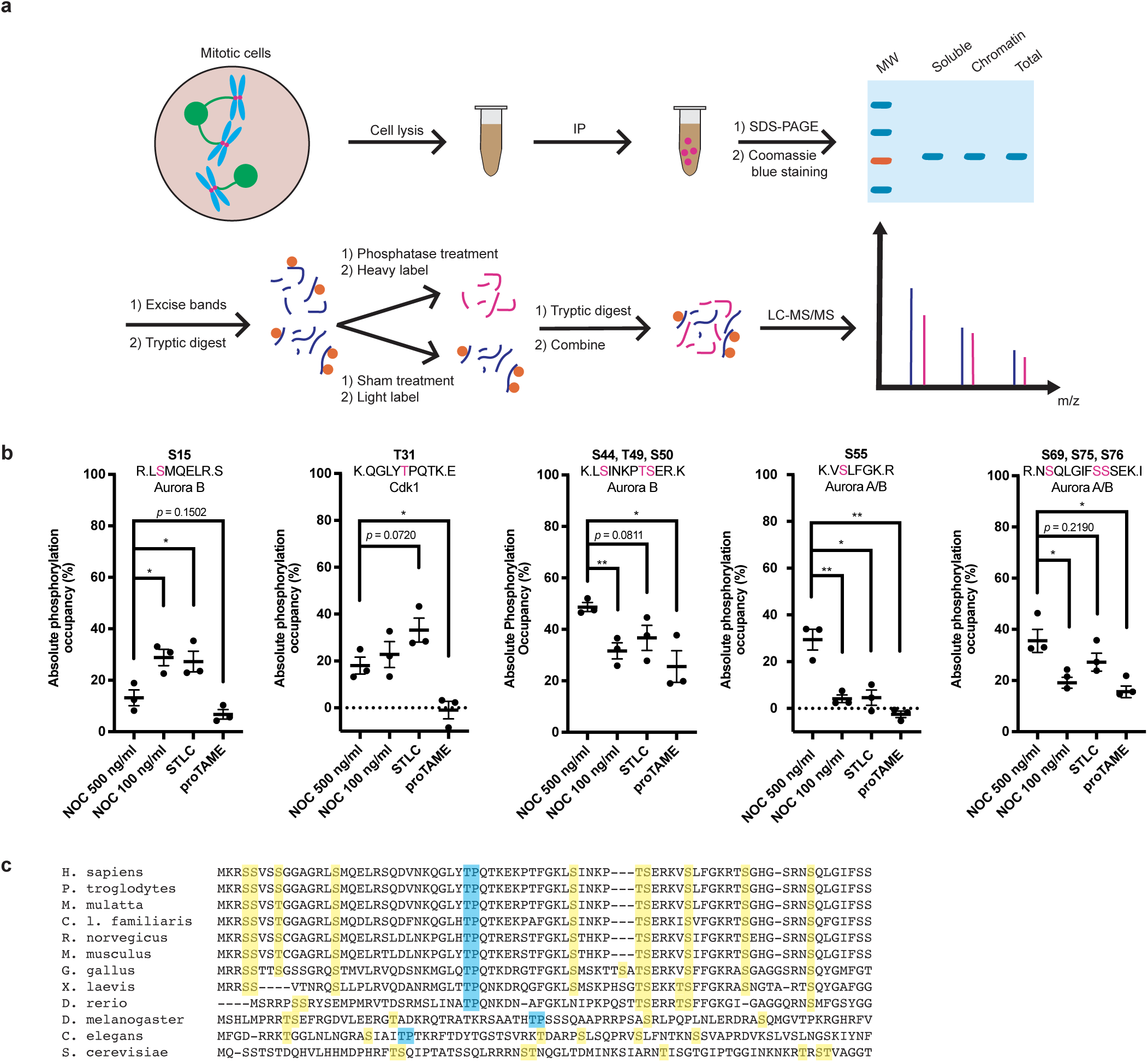
Hec1 is phosphorylated at only partial occupancy during mitosis. A) Cartoon illustration depicting the experimental scheme used to determine protein phosphorylation occupancy. Mitotic cells are lysed. Hec1 is then immmunoprecipitated from the lysate and purified by SDS-PAGE. The band of Hec1 is then excised, and the protein subjected to tryptic digest. The sample is then divided in two, and half treated with CIP and labelled with a heavy isotope. The sample is then re-combined and analyzed by MS. B) Histograms showing the absolute phosphorylation occupancy of various Hec1 peptides purified from total cell lysate as determined by MS. The individual and average values from 3 independent experiments are shown. Error bars indicate mean +/-SEM. * denotes p < 0.05, ** denotes p < 0.01. C) Amino acid sequence alignment of Hec1 orthologs from residues 1-76. Residues highlighted in yellow indicate Aurora kinase consensus sites. Residues highlighted in blue indicate Cdk1 consensus sites.

We obtained sequence coverage and determined the phosphorylation occupancy of five known phosphorylation sites within the n-terminal tail of Hec1: S15, S44, T49/S50, S55 and S69, as well as the previously identified, but uncharacterized phosphorylation sites, T31, S75 and S76 ^20^ (Figure 1B). We were unable to determine the occupancy of the known phosphorylation sites S4, S5, S8 and S62 because their peptides are very hydrophilic and appeared only sporadically in the MS data. The Aurora B phosphorylation site S15 and the Aurora A/B phosphorylation site S55 were confidently localized on separate peptides for which occupancy could be determined. Multiple peptides spanning the Aurora B phosphoacceptors S44, T49 and S50 were identified as singly phosphorylated at S44 and separately at either T49 or S50, which could not be confidently localized; in such cases, it is reasonable to assume that either site could be phosphorylated, as determined previously ^20–22^. We did not observe any peptides doubly phosphorylated on S44 and S49/T50. Furthermore, singly phosphorylated peptides containing the Aurora A and/or B phosphoacceptors S69, S75 and S76 were observed in which S69 phosphorylation was unambiguously assigned on some peptides, as well as on either S75 or S76, which has been previously observed ^20^. S75/S76 phosphorylation has been shown to decrease under Aurora B inhibition ^20^, although the adjacent amino acid sequence is not characteristic of mitotic kinases ^23^, suggesting that it may be an atypical Aurora B consensus motif. Finally, we could determine phospho-occupancy on a single peptide containing T31, a site that is conserved and conforms to a consensus sequence typical of Cdk1 phosphorylation sites ^23^ (Figure 1C).

The direction and magnitude of the phospho-occupancy changes of each peptide among the four conditions that we examined revealed three categories of response (Figure 1B). One category includes S44/T49/S50 and S69/S75/S76, which displayed maximum occupancy in mitotic cells without microtubule attachments (500 ng/ml NOC) and with approximately 2-fold decrease of phospho-occupancy in each of the other conditions when microtubules are present (irrespective of the attachment configuration). Another category is defined by the peptides containing S15 and T31 that are Aurora B and Cdk1 sites, respectively. These sites display maximum phospho-occupancy under conditions that generate mostly erroneous k-MT attachments (STLC and 100 ng/ml NOC), and exceptionally low occupancy in cells with bi-oriented k-MT attachments (proTAME). A third category is defined by the peptide containing S55, which displayed switch-like behavior with maximum phosphorylation in the absence of microtubules (500 ng/ml NOC) and almost no phosphorylation in the other conditions. These findings are broadly consistent with previous experiments showing that Hec1 phosphorylation is lowest in mitotic cells with bi-oriented k-MT attachments ^5^, but demonstrates that these sites respond independently to different types of k-MT attachments. Furthermore, the phospho-occupancy of individual sites does not uniformly correlate with inter-kinetochore distance that has previously been proposed to influence the extent of Hec1 phosphorylation. For example, the S15 and S44/T49/S50 sites have substantially different phospho-occupancies under conditions with similar average inter-kinetochore distances (500 ng/ml vs 100 ng/ml NOC) (Figure 1B, S2B).

Mutational and domain swapping experiments suggest that the Hec1 n-terminal tail phosphorylation sites participate interchangeably in causing detachment of microtubules from Hec1 ^8^. Therefore, the functional detachment of microtubules is likely sensitive to the cumulative phosphorylation occupancy of all these sites, and our data can express the phospho-occupancies within this domain as a cumulative score. The theoretical maximum cumulative occupancy for the Hec1 peptides as measured by MS is 5.00 (five peptides, each with 100% occupancy). Surprisingly, in any of the four conditions that we examined, with a range of k-MT attachments, the cumulative phosphorylation of all peptides did not approach this maximum, and none of the individual peptides that we sampled exceeded 50% occupancy. The condition displaying the highest total occupancy was mitotic cells treated with 500 ng/ml of NOC, which equalled 1.46, or approximately 29.2% of theoretical maximum occupancy. Cells treated with STLC and 100 ng/ml NOC that have microtubule attachments, albeit mostly erroneous ones, displayed a cumulative occupancy of 1.37 (27.4% of theoretical maximum) and 1.08 (21.6% of theoretical maximum), respectively. Cells under proTAME arrest displayed a cumulative occupancy of 0.46 (9.2% of theoretical maximum). Similar trends were observed on Hec1 purified from soluble or chromatin-associated cell fractions (S2E), indicating that phospho-occupancy is not dependent on specific protein subpopulations. Thus, the cumulative occupancy of these Hec1 peptides changes by only 20% (from approximately 29% to 9%) between mitotic cells with no k-MT attachments and those with bi-oriented k-MT attachments.

### Hec1 phosphorylation on T31 during mitosis is temporally regulated

To investigate how Hec1 phospho-occupancy is set by the competing activities of protein kinases and phosphatases, we generated an antibody against phosphorylated T31 (pT31). We focused our analysis on T31 as it is the only predicted Cdk1 site in the n-terminal tail of Hec1 and it is uncharacterized. We used immunofluorescence (IF) microscopy to validate its specificity towards pT31 in mitotic cells and not to other phosphorylated sites within the Hec1 tail (Figure S3A). Importantly, the relative pT31 intensity at kinetochores in imaged cells paralleled the occupancy values determined using MS in that T31 is poorly phosphorylated in cells arrested in 500 ng/ml NOC or proTAME, but robustly in cells arrested in 100 ng/ml NOC or STLC (Figure 1B, S3D, S3E).

To explore the functional role of T31 phosphorylation, we measured the pT31 intensity at kinetochores in human cells as they underwent an unperturbed mitosis. We found that in both HeLa and non-transformed RPE1 cells, there is little pT31 in prophase, the intensity sharply increases in prometaphase and then declines in metaphase and anaphase (Figure 2A, 2B, S3B, S3C). We also measured the pT31 levels by IF in cells arrested under the same conditions that we used for MS analysis. Since the cells were treated identically for both IF and MS, the phospho-occupancy values determined by MS can be used to calibrate the fluorescence intensities at kinetochores in cells progressing through mitosis. Specifically, the fluorescence intensity at kinetochores in cells arrested in STLC would reflect approximately 33% T31 phospho-occupancy. Using this calibration, the pT31 levels range from <10% (prophase) to ∼40% (late prometaphase) (Figure 2A, 2B).

**Figure 2:**
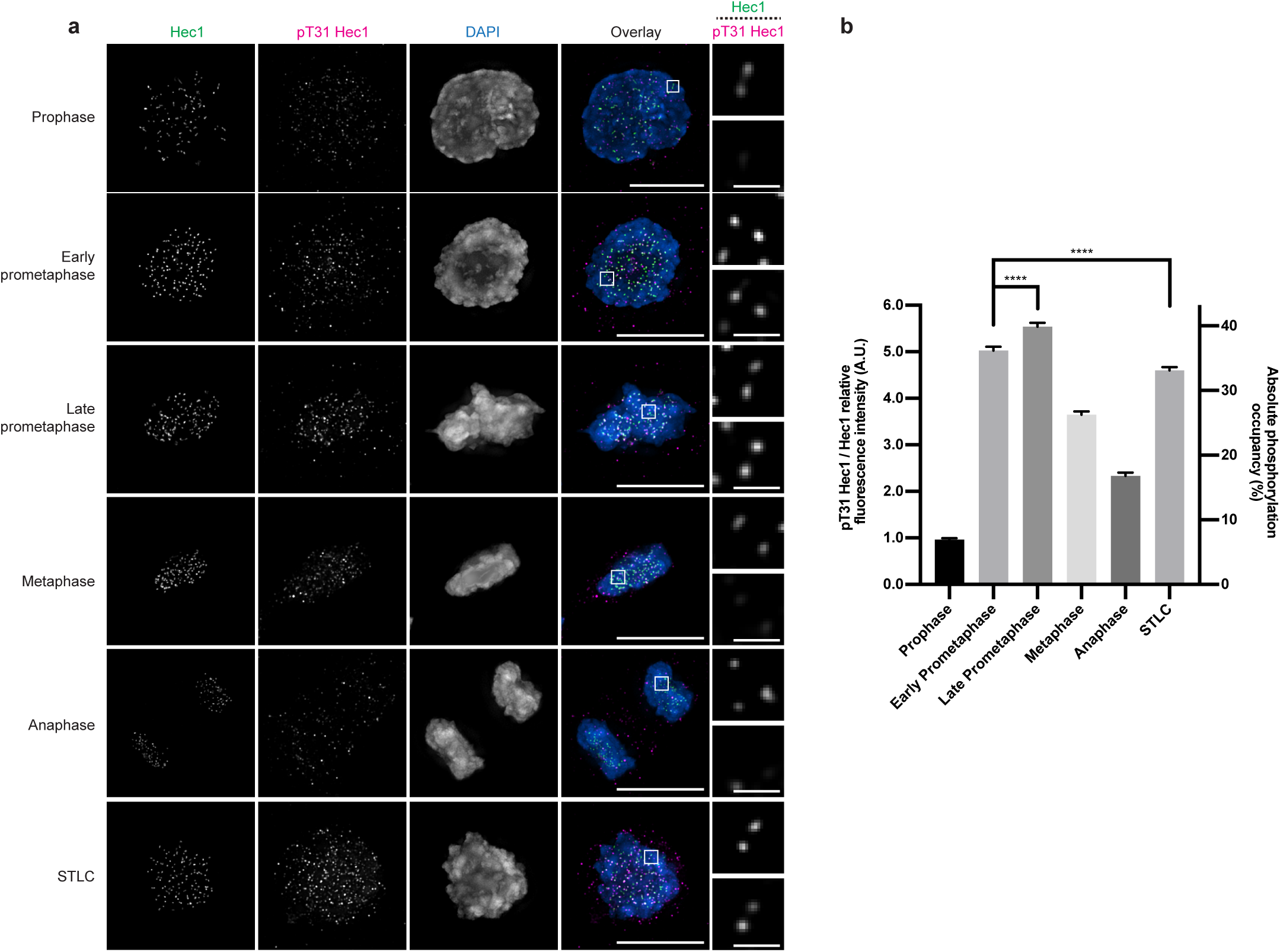
Hec1 phosphorylation on T31 is temporally regulated during mitosis. A) Immunofluorescence images from an asynchronous population of HeLa cells in various stages of mitosis or arrested with STLC stained for Hec1, pT31 Hec1 and DAPI. The Hec1 and pT31 channels were adjusted evenly for brightness and contrast for presentation. The DAPI channel of each condition was adjusted independently. Representative images from 3 independent experiments are shown. The scale bars of main images are 10 μm, and insets are 1 μm. B) Quantification of the relative kinetochore intensities from the conditions from panel (A). The condition with the lowest level of pT31 Hec1/Hec1 was set to 1 and the other conditions shown as fold-changes. 25 kinetochores were quantified from each of 20 cells for each of 3 independent repeats. Error bars indicate the mean +/- SEM. The source data and images for the STLC condition originate from the experiment shown in Extended data 3c, 3f.

### Hec1 is phosphorylated on T31 by Cdk1-Cyclin B1 complexes

Human cells enter mitosis with Cdk1-Cyclin B1 and Cdk1-Cyclin A2 complexes which phosphorylate residues at the same consensus motif ^23^. We therefore asked which of these kinases phosphorylates Hec1 on T31 by siRNA mediated gene silencing. Strikingly, T31 phosphorylation was abolished at kinetochores of mitotic cells lacking either Cyclin A2 or Cyclin B1 (Figure 3A, 3B). However, we also observed the failure of Cyclin B1-GFP to localize to kinetochores in cells lacking Cyclin A2 (Figure 3C). These results indicate that Cyclin A2 is an upstream factor required for Cyclin B1 kinetochore localization and that Cdk1-Cyclin B1 is likely to be directly responsible for Hec1 T31 phosphorylation.

**Figure 3:**
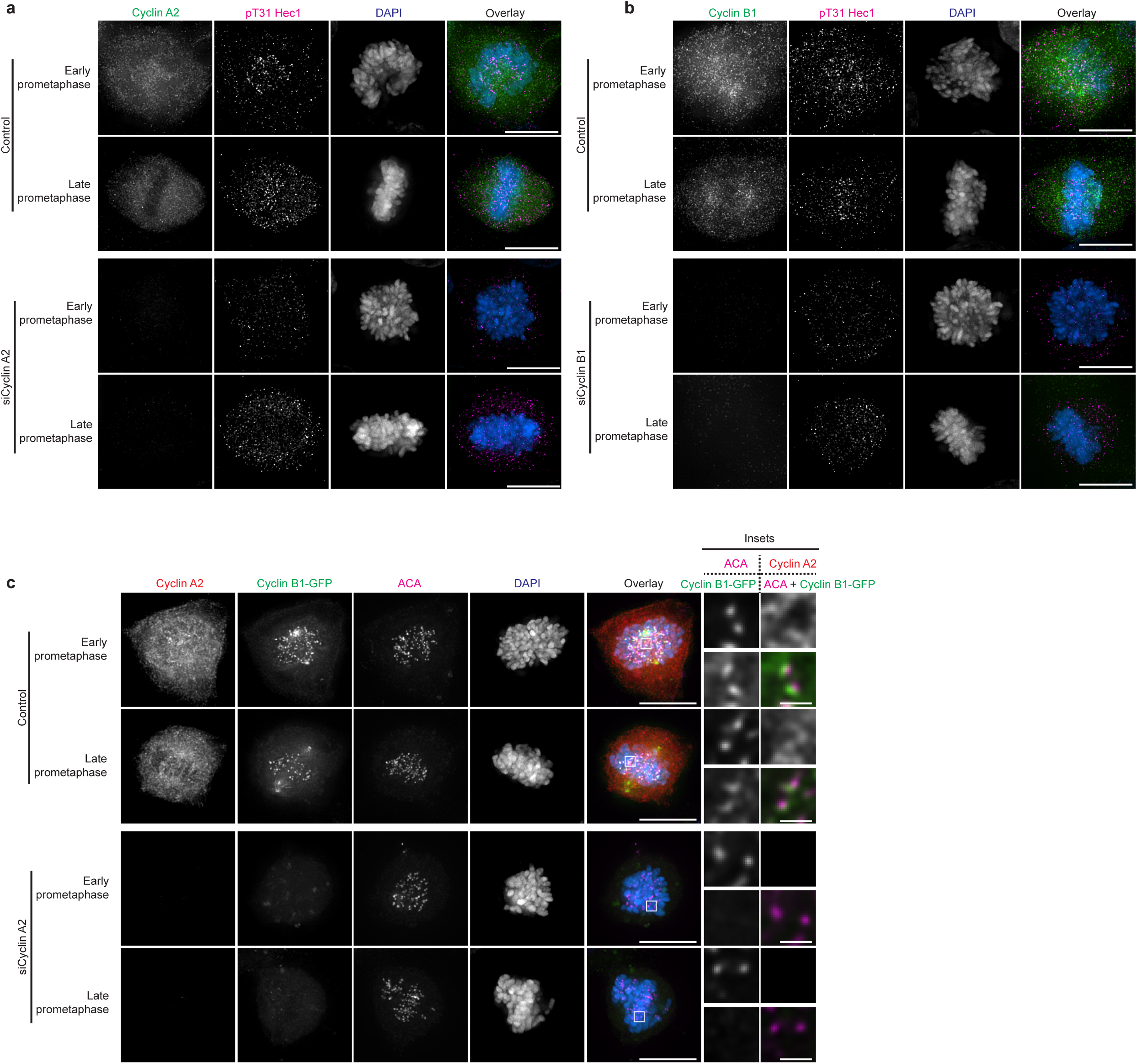
Both Cyclin A2 and B1 are required for Hec1 T31 phosphorylation; Cyclin A2 is required for kinetochore localization of Cyclin B1 during prometaphase. A) Immunofluorescence images from an asynchronous population of HeLa cells transfected with control or siRNA against Cyclin A2 and then stained for Cyclin A2, pT31 Hec1 and DAPI. The Cyclin A2 and Hec1 pT31 channels were adjusted evenly for brightness and contrast. The DAPI channel was adjusted independently. The scale bars are 10 μm. Representative images of 3 independent experiments are shown. B) Immunofluorescence images from an asynchronous population of HeLa cells transfected with control or siRNA against Cyclin B1 and then stained for Cyclin B1, pT31 Hec1 and DAPI. The Cyclin B1 and Hec1 pT31 channels were adjusted evenly for brightness and contrast. The DAPI channel was adjusted independently. The scale bars are 10 μm. Representative images of 3 independent experiments are shown. C) Immunofluorescence images from an asynchronous population of HeLa Cyclin B1-GFP cells transfected with control or siRNA against Cyclin A2 and then stained for Cyclin A2, ACA and DAPI. The GFP, Cyclin A2, and ACA channels were adjusted evenly for brightness and contrast. The DAPI channel was adjusted independently. The scale bars for the main images are 10 μm and 1 μm for the insets. Representative images of 2 independent experiments are shown.

Next, we assessed the localization of Cyclins A2 and B1 relative to Hec1 and pT31. Consistently ^24–27^, Cyclin A2-GFP is cytosolic and does not obviously localize to kinetochores (Figure 4A, 4B), while Cyclin B1-GFP localized to kinetochores, microtubules and spindle poles in early through mid-prometaphase and then progressively disappeared as cells reached metaphase. Kinetochore-localized Cyclin B1-GFP largely co-localized with Hec1 and pT31, although it occupies a larger volume than Hec1 (Figure 4C-E).

**Figure 4:**
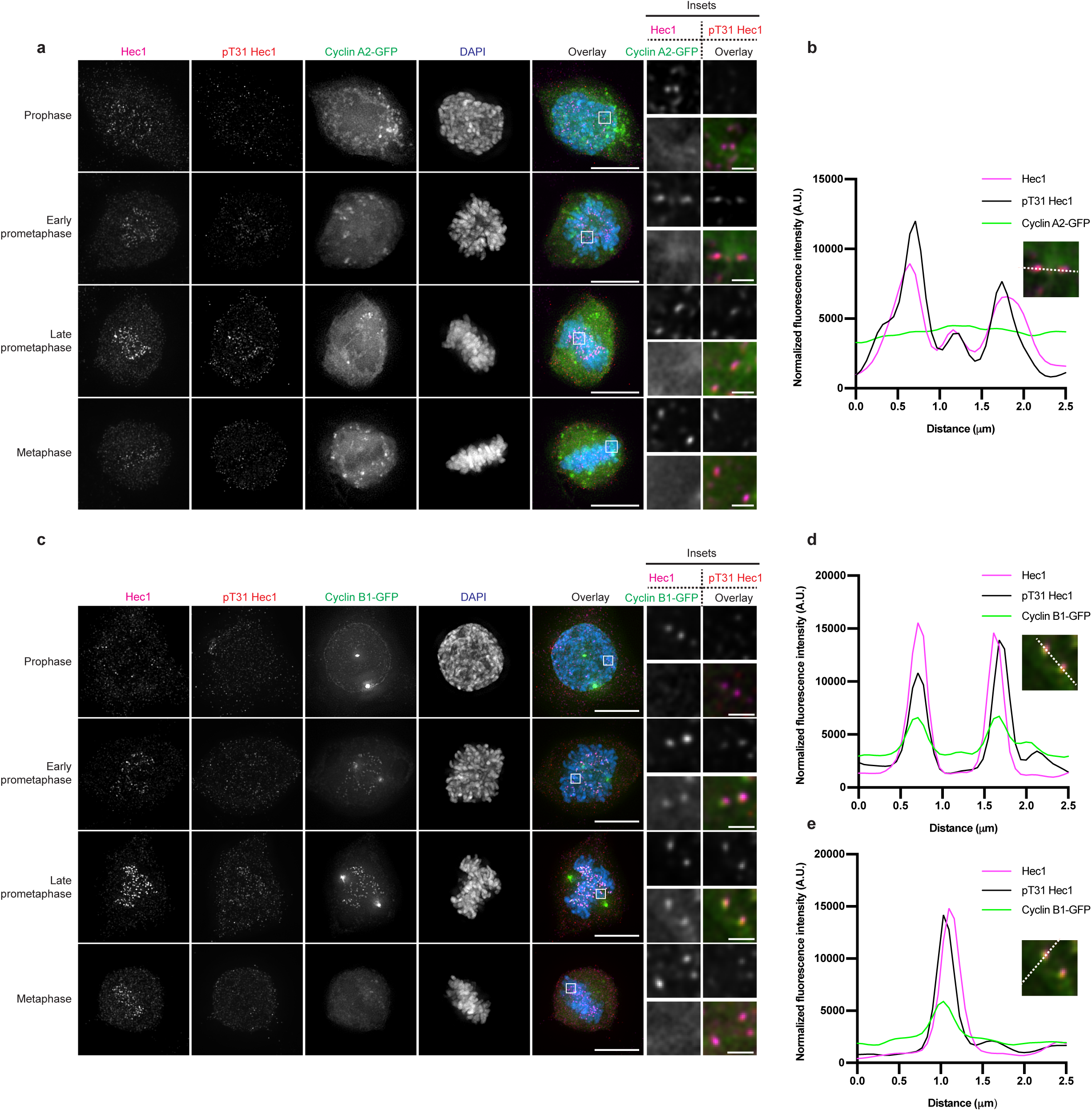
Cyclin B1 but not Cyclin A2 co-localizes with Hec1 during prometaphase. A) Immunofluorescence images from an asynchronous population of HeLa Cyclin A2-GFP cells stained for Hec1, pT31 Hec1 and DAPI. The Hec1 and Hec1 pT31 channels were adjusted evenly for brightness and contrast. The GFP and DAPI channels were adjusted independently. The scale bars for the main images are 10 μm and 1 μm for the insets. Representative images of 2 independent experiments are shown. B) Line scan through the indicated pair of kinetochores from the cells in panel (A). C) Immunofluorescence images from an asynchronous population of HeLa Cyclin B1-GFP cells stained for Hec1, pT31 Hec1 and DAPI. The Hec1 and Hec1 pT31 channels were adjusted evenly for brightness and contrast. The GFP and DAPI channels were adjusted independently. The scale bars for the main images are 10 μm and 1 μm for the insets. Representative images of 2 independent experiments are shown. D) Line scan through the indicated pair of kinetochores from the cells in panel (C.) E) Orthogonal line scan through the upper kinetochore from the cells in panel (C).

### Hec1 T31 phosphorylation is increased on kinetochores with improper microtubule attachments

The T31 location within the amino acid sequence of Hec1 and the temporal profile of pT31 suggest a role for pT31 in k-MT attachment error correction. To explore this idea, we first measured the pT31 intensity at laterally attached compared to end-on attached kinetochores (Figure 5A, 5B). This experiment revealed an approximately 2-fold increase in pT31 at laterally attached compared to end-on attached kinetochores. We then purposefully induced k-MT attachment errors using the CENP-E inhibitor GSK-923295, which induces clustering of chromosomes at the spindle poles and k-MT attachment errors through persistent monotelic attachments ^28, 29^. This treatment induced an approximately 2-fold increase in pT31 at pole-arrested compared to aligned kinetochores (Figure S5A, S5B). As an additional but different means of inducing errors in k-MT attachments, we utilized a mutant of Hec1 (Hec1 8A) ^6, 30^ (Figure S7A) which prevents complete chromosome alignment through hyperstable k-MT attachments that cannot be corrected. In cells expressing Hec1 8A, a majority of chromosomes eventually align with bi-orientated attachments, but some remain unaligned (Figure S5C). Inter-kinetochore distance measurements confirm hyperstabilization of k-MT attachments in mitotic cells expressing Hec1 8A (Figure S7B) ^5, 14, 31, 32^. In these cells, pT31 levels were also elevated in pole-arrested compared to aligned kinetochores (Figure S5C, S5D). Taken together, these data show increased phospho-occupancy at T31 on improper lateral and monotelic attachment configurations compared to end-on k-MT attachments.

**Figure 5:**
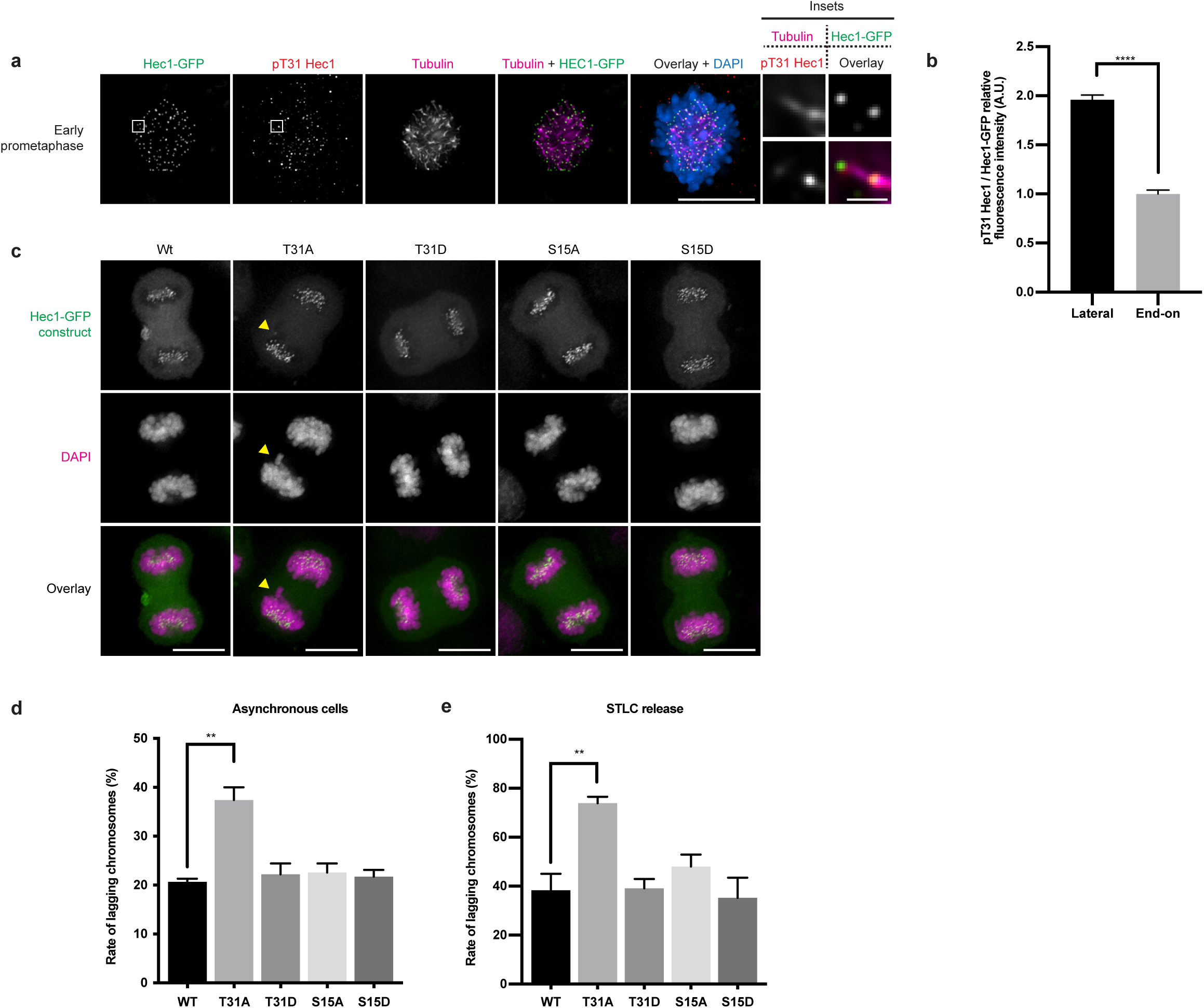
Hec1 T31 phosphorylation is critical for k-MT attachment error correction. A) Immunofluorescence images of HeLa doxycycline inducible Hec1 knockout cells transfected with wild-type Hec1-GFP in an early stage of prometaphase stained for pT31 Hec1, tubulin and DAPI. The images were adjusted for brightness and contrast for presentation. Representative images from 2 independent experiments are shown. The scale bars of main images are 10 μm, and in the insets are 1 μm. B) Quantification of the relative kinetochore intensities of end-on attached kinetochores compared to laterally attached kinetochores as shown in panel (A). The levels of the end-on attached kinetochores were set to 1 and the levels of the laterally attached kinetochores shown as a fold-change. 827 (laterally attached) and 505 (end-on attached) total kinetochores from 10 cells for each of 2 independent experiments were measured. Error bars indicate the mean +/-SEM. **** denotes p < 0.0001. C) Immunofluorescence images of HeLa doxycycline inducible Hec1 knockout cells transfected with the indicated Hec1-GFP constructs. 72 hours post transfection the cells were fixed and stained with DAPI. The GFP and DAPI channels were adjusted evenly for brightness and contrast for presentation. A lagging chromosome is indicated with a yellow arrow. Representative images from 3 independent experiments are shown. The scale bars are 10 μm. D) Quantification of the percentage of lagging chromosomes in cells from an asynchronous population undergoing anaphase and expressing the indicated Hec1 constructs. At least 319 cells per construct were scored for the presence of lagging chromosomes over 3 independent repeats. Error bars indicate the mean +/- SEM. ** denotes p < 0.01. E) Quantification of the percentage of lagging chromosomes in anaphase for cells expressing the indicated Hec1 constructs following an STLC washout. 120 cells per construct were scored for the presence of lagging chromosomes for each of 3 independent repeats. Error bars indicate the mean +/- SEM. ** denotes p < 0.01.

To test if Cyclin B1 specifically localizes to kinetochores with improper k-MT attachments, we treated cells with GSK-923295 to generate k-MT attachment errors. We frequently observed pole-arrested kinetochores with lateral monotelic attachments where only the attached kinetochore showed Cyclin B1 present. We also observed kinetochores with lateral syntelic attachments with Cyclin B1 present on both sister kinetochores. We did not observe any Cyclin B1 on kinetochores with end-on amphitelic attachments (Figure S5E, S5F). Furthermore, the intensity of Cyclin B1-GFP at kinetochores was high in mitotic cells arrested with 100 ng/ml NOC or STLC, and low in cells arrested with 500 ng/ml NOC or proTAME (Figure S4A, S4B), a pattern similar to pT31 levels in cells arrested under those conditions (Figure S3D, S3E). Thus, Cyclin B1 localization is enriched on improper k-MT attachments coinciding with increased Hec1 pT31.

Persistent errors in k-MT attachment manifest as chromosomes lagging in anaphase and increase chromosome mis-segregation ^33^. To test if Hec1 T31 phosphorylation participates in error correction, we expressed Hec1-GFP in which T31 was mutated to either alanine (T31A) or aspartic acid (T31D). For comparison, we also generated S15A and S15D mutants because S15 is phosphorylated by a different kinase (Aurora B) and it demonstrated a similar occupancy profile as T31 (Figure 1B, S2E). Hec1 T31A expression resulted in elevated lagging chromosome rates in asynchronous cells (Figure 5C, 5D). Hec1 T31D expression showed no change relative to wild-type Hec1, suggesting that this substitution may not have a functional effect. Next, we increased k-MT attachment errors via treatment of cells with an STLC washout strategy and measured the lagging chromosome frequency. As expected, the overall lagging chromosome rate was elevated in all cases. Hec1 T31A expression resulted in very high lagging chromosome rates in cells recovering from STLC treatment, suggesting that Hec1 T31 phosphorylation is required for efficient k-MT attachment error correction (Figure 5E). Interestingly, Hec1 S15A or S15D expression had no significant effect on lagging chromosomes rates in either asynchronous cells or cells recovering from STLC treatment (Figure 5C-E), suggesting perhaps that Aurora kinases can compensate for the absence of one phosphorylation site by increasing phosphorylation at other sites. These data demonstrate that Hec1 T31 phosphorylation by Cdk1-Cyclin B1 is important for efficient k-MT attachment error correction and that pT31 provides a greater contribution to error correction than Hec1 S15 phosphorylation by Aurora B.

### Kinetochore Hec1 phosphorylation levels are determined by phosphatase activity

To determine the phosphatase contribution to the net phospho-occupancy of the Cdk1-Cyclin B1 site T31, we arrested cells in mitosis with 100 or 500 ng/ml NOC and then treated the cells with the general phosphatase inhibitor okadaic acid (OA; 200 nM) for 2 hours prior to fixation. In both conditions, the pT31 intensity at kinetochores increased in the presence of okadaic acid, but not to saturation (Figure 6A, 6B). This occurred in spite of poor localization of Cyclin B1 to kinetochores lacking microtubule attachments (500 ng/ml NOC) suggesting that even partial localization of Cyclin B1 is sufficient to elevate Hec1 phospho-occupancy under these conditions. We also tested the contribution of phosphatases to the levels of pS44 in cells under the same conditions. Interestingly, the extent of phosphorylation was markedly increased in cells treated with the high dose of nocodazole, but only to a lesser extent in cells treated arrested in the lower dose (Figure 6C, 6D). These data show that the phospho-occupancy of Aurora and Cdk1 substrates are differentially regulated. We also noticed in these experiments that in cells treated with 100 ng/ml NOC the kinetochores were clustered around and attached to centrosomes with short microtubule stub-spindles (Figure 6A, 6C, S2A, S3D, S4A). However, following treatment with OA, the kinetochores were distributed throughout the cytoplasm, suggesting that the kinetochores had detached from the microtubules. To determine if this was the case, we stained identically treated cells for α tubulin and examined the extent of kinetochore attachment. This experiment confirmed that in cells treated with nocodazole only, all chromosomes were attached to the stub spindles by at least one kinetochore. However, in cells treated with both nocodazole and OA almost all kinetochores were detached from the spindles (Figure 6E), suggesting that Hec1 phosphorylation levels are repressed by phosphatases to prevent k-MT detachment despite ongoing kinase activity. Given that only a modest 1.3-fold increase in phosphorylation was observed at S44 in cells treated with OA, k-MT attachments are likely very sensitive to phosphorylation levels.

**Figure 6:**
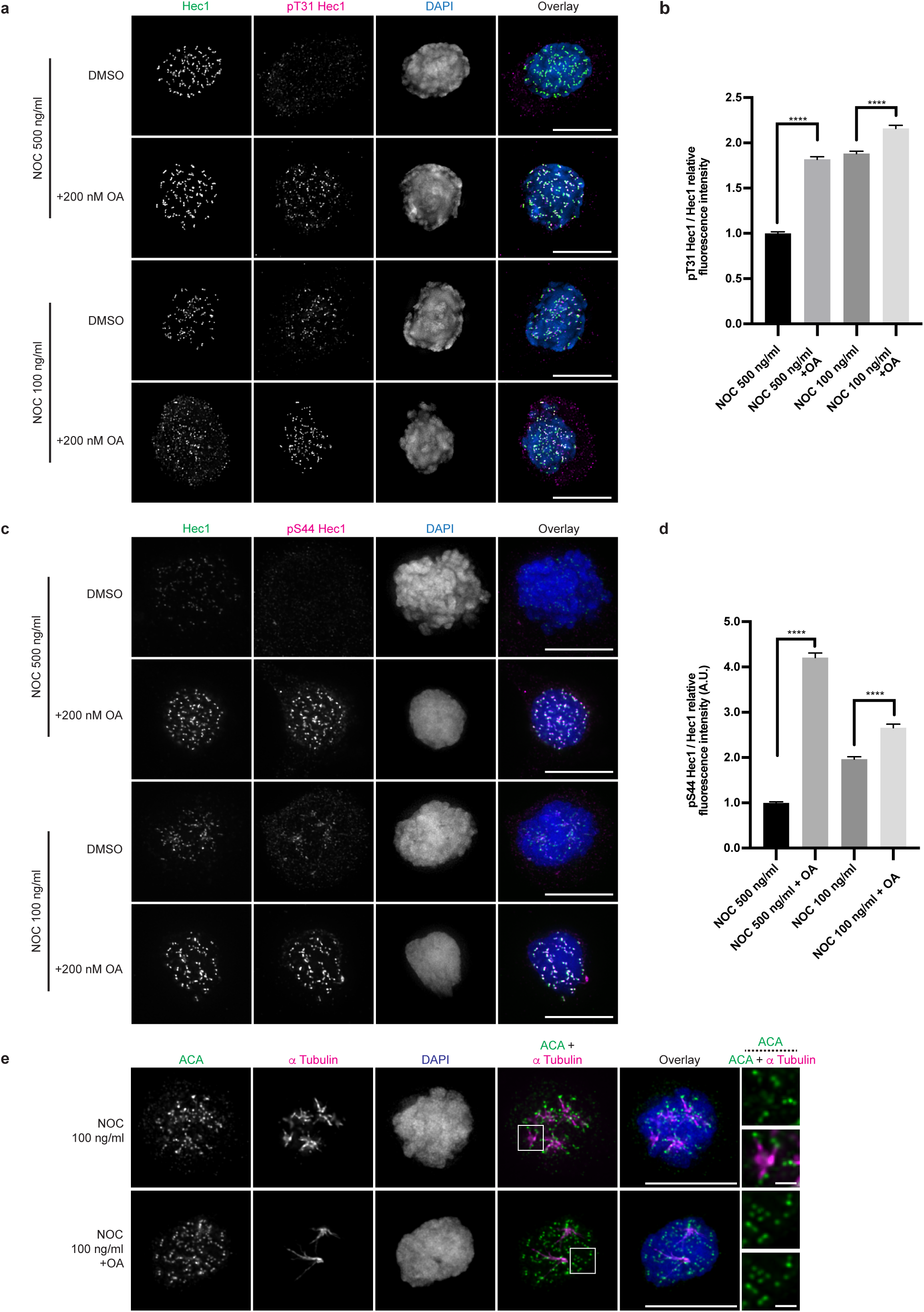
Hec1 phosphorylation is repressed by phosphatase activity to bias kinetochores towards a state of attachment. A) Immunofluorescence images of HeLa cells synchronized by thymidine and then NOC as indicated. The cells were then treated for 2 hours with OA or DMSO control and finally fixed and stained for Hec1, pT31 Hec1 and DAPI. The Hec1 and pT31 Hec1 channels were adjusted evenly for brightness and contrast. The DAPI channel was adjusted independently. The scale bars are 10 μm. Representative images of 3 independent experiments are shown. B) Quantification of the relative kinetochore intensities from the conditions in panel (A). The condition with the lowest level of pT31 was set to 1 and the other conditions shown as fold-changes. 25 kinetochores were quantified from each of 20 cells for each of 3 independent repeats. Error bars indicate the mean +/- SEM. **** denotes p < 0.0001. C) Immunofluorescence images of cells prepared as in panel (A), but stained for pS44 instead of pT31. D) Quantification of the relative kinetochore intensities from the conditions in panel (C). The condition with the lowest level of pS44 was set to 1 and the other conditions shown as fold-changes. 25 kinetochores were quantified from each of 20 cells for each of 3 independent repeats. Error bars indicate the mean +/- SEM. **** denotes p < 0.0001 E) Immunofluorescence images of HeLa cells synchronized by thymidine and then 100ng / ml NOC as indicated. The cells were then treated for 2 hours with OA or DMSO control and finally fixed and stained for ACA and α Tubulin. The scale bars for main images are 10 μm and 1 μm for insets.

K-MT attachments have previously been shown to be regulated during prometaphase by the PP2A-B56 family of phosphatases ^10, 11^. To determine if Hec1 tail phosphorylation is regulated by PP2A-B56, we took advantage of the fact that kinetochore isoforms of B56 are recruited to the kinetochore by the protein BubR1 ^11^. Therefore, knockdown of BubR1 removes them from the kinetochore. We silenced expression of BubR1 using siRNA mediated knockdown (Figure S6K) and examined the phosphorylation state of Hec1. Interestingly, we observed an approximately 1.2-fold increase in phosphorylation of both S44 and T31 positions on Hec1 (Figure S6A-D), which suggests that Hec1 phosphorylation by both Aurora and Cdk1 kinases are repressed by the PP2A-B56 group of phosphatases and is consistent with previous findings ^10, 11^. In these experiments we also observed a large proportion of the BubR1 knockdown cells that displayed chromosome alignment defects (Figure S6A, S6C, S6E, S6I) as reported previously ^11^. Since it had been previously determined that the alignment defects are due to a loss of k-MT detachment as a result of increased KMN network protein phosphorylation ^10, 11^, we also examined the stability of MTs using a cold MT stability assay under these conditions. Despite an only 1.2-fold increase in phosphorylation at S44 and T31 on Hec1, this experiment revealed a 0.4-fold loss of MTs (Figure S6E, S6F), again suggesting that the k-MT attachment state is hypersensitive to KMN network protein phosphorylation.

To confirm that Hec1 phosphorylation is repressed by PP2A-B56, we silenced the expression of the B56 subunit isoforms using a previously described pool of siRNAs ^10^ (Figure S6L) and examined the phosphorylation state of Hec1 at S44 and T31. This experiment showed a 1.6-fold increase in pS44 levels (Figure 7A, 7B). Interestingly, under the same conditions we observed a slight decrease in pT31 levels (Figure 7C, 7D), although we continued to observe increased phosphorylation at pole-arrested kinetochores compared to aligned kinetochores (Figure 7C). This might indicate additional OA-sensitive phosphatases acting to regulate the phospho-occupancy of the T31 site. Consistent with previous findings ^10^, we also observed that cells with B56 knockdown display moderate to severe chromosome alignment defects (Figure 7A, 7C, 7E, S6G). To test if the defects in chromosome alignment are due to a loss of K-MT attachment stability, we again performed a cold MT stability assay in cells with B56 knockdown compared to controls. This experiment revealed that cells with B56 knockdown have only 35% of the cold-stable microtubule fluorescence compared to control cells, which likely explains the defects in chromosome alignment. As we and others have noted previously, the presence of (improper) k-MT attachments is required both for Cyclin B1 localization to kinetochores and for robust phosphorylation of T31 (Figure S4) ^26, 27^. We therefore examined if kinetochores have such weak attachments under conditions of total phosphatase loss that Cdk1-Cyclin B1 is unable to efficiently localize to and phosphorylate Hec1 under conditions of phosphatase loss at kinetochores. We compared the localization pattern of Cyclin B1-GFP during prometaphase under conditions of BubR1 knockdown and also B56 knockdown. Interestingly, most cells with BubR1 knockdown maintained Cyclin B1 localization at kinetochores, as did cells with B56 knockdown and moderate chromosome alignment defects. However, most cells with B56 knockdown and severe alignment defects did not have Cyclin B1 at kinetochores (Figure S6G-J). This suggests that Cyclin B1 might only localize to kinetochores that are strongly bound to microtubules in order to destabilize them, whereas it would not further destabilize attachments that are already weak.

**Figure 7:**
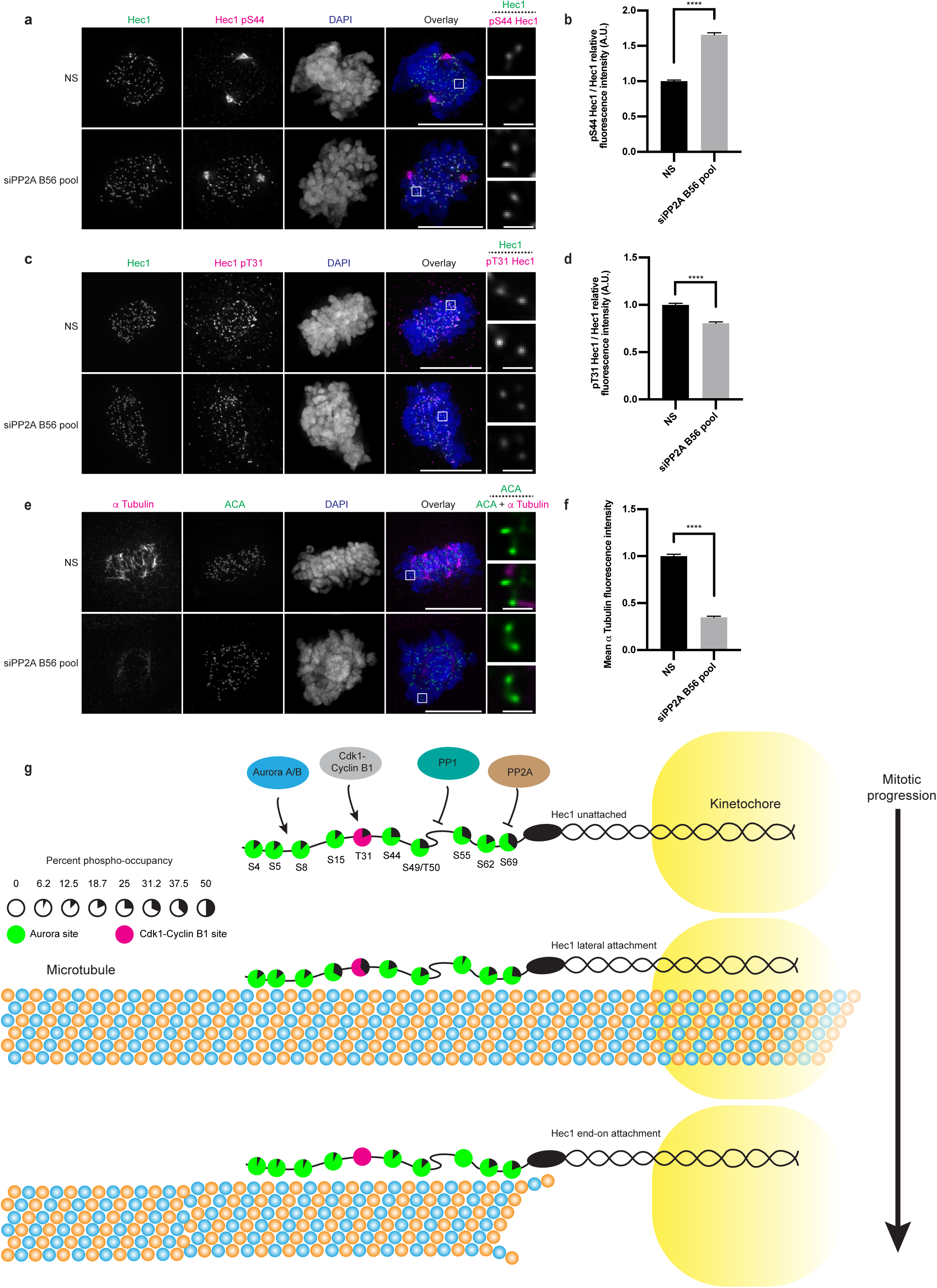
Hec1 phosphorylation is repressed by PP2A-B56 with differential effects on pT31 and pS44. A) Immunofluorescence images from an asynchronous population of HeLa cells transfected with control or a pool of siRNA against B56 α-ε and then stained for Hec1 pS44, Hec1 and DAPI. The Hec1 and Hec1 pS44 channels were adjusted evenly for brightness and contrast. The DAPI channel was adjusted independently. The scale bars for main images are 10 μm and 1 μm for insets. Representative images of 3 independent experiments are shown. B) Quantification of the relative kinetochore intensities from the conditions in panel (A). The condition with the lowest level of pS44 was set to 1 and the other conditions shown as fold-changes. 25 kinetochores were quantified from each of 20 cells for each of 3 independent repeats. Error bars indicate the mean +/- SEM. **** denotes p < 0.0001. C) Immunofluorescence images of cells prepared as in panel (A) except stained for pT31 instead of pS44. D) Quantification of the relative kinetochore intensities from the conditions in panel (C). The condition with the lowest level of pT31 was set to 1 and the other conditions shown as fold-changes. 25 kinetochores were quantified from each of 20 cells for each of 3 independent repeats. Error bars indicate the mean +/- SEM. **** denotes p < 0.0001. E) Immunofluorescence images from an asynchronous population of HeLa cells transfected with control or a pool of siRNA against B56 α-ε. The media was then changed for media at 4°C, and the cells placed at 4°C for 15 minutes prior to fixation and staining with antibodies against ACA and α Tubulin. Representative images of 3 independent experiments are shown. F) Quantification of the relative intensity of α Tubulin. The levels of the control condition were set to 1 and the other condition shown as a fold-change. At least 50 cells per condition from each of 3 independent experiments were quantified. Error bars indicate the mean +/- SEM. **** denotes p < 0.0001. G) Cartoon illustration showing the kinase and phosphatase network regulating Hec1 phosphorylation. In early prometaphase, cells without k-MT attachments have elevated levels of fractional phosphorytion on several Hec1 sites that are phosphorylated by Aurora kinases (green circles) or Cdk1-Cyclin B1 kinase (magenta circles). When lateral k-MT attachments occur, the levels of phosphorylation generally decrease. Finally, when end-on k-MT attachments occur, levels of phosphorylation decrease further. Phosphorylation levels at Hec1 S4,5,8 and S62 are speculative or based on partial data coverage.

### The Aurora and Cdk1 phosphorylation networks respond differentially to stimuli

We next sought to compare how the Aurora B and CDK1 signaling networks respond to alterations in k-MT stability. We used the Hec1 8A and 8D mutants (Figure S7A) because they preserve an Aurora B site (S44) and the Cdk1 site (T31), allowing us to monitor the phosphorylation status of these two specific sites in cells with k-MT attachment errors. We observed a significant increase in pS44 levels at kinetochores in early and late prometaphase cells expressing Hec1 8A compared to wild-type Hec1. We also observed a slight increase in pS44 levels in late prometaphase cells expressing the 8D mutant (Figure S7C, S7E). In contrast, there was a significant decrease in pT31 levels in early prometaphase cells expressing Hec1 8A (Figure S7D, S7F). Since this Hec1 mutant promotes the formation of precocious end-on k-MT attachments in early prometaphase ^32^, this is consistent with our results showing decreased pT31 at end-on attached kinetochores (Figure 5A, 5B). There was no overall significant change in pT31 levels in late prometaphase Hec1 8A expressing cells (although we observe increased phosphorylation at pole-arrested kinetochores), suggesting a temporal change in Cdk1 or phosphatase activity for site T31 under these conditions. Notably, we observed a slight decrease in pT31 levels in cells expressing the 8D mutant in both early and late prometaphase cells (Figure S7D, S7F) which is consistent with our data showing that phosphatase inhibition reduces pT31 levels due to a loss of k-MT attachment and Cyclin B1 localization at kinetochores (Figure 7, S6G, S6H). Thus, the Aurora B and Cdk1 signaling networks respond independently to changes in the nature of k-MT attachments, which is consistent with our phospho-occupancy measurements determined by MS (Figure 1B).

## Discussion

Taken together, our data demonstrate that phosphorylation of the population of Hec1 molecules that form an essential component of the kinetochore-microtubule attachment interface does not follow an all-or-none mechanism despite the fact that phosphorylation of individual amino acids on individual molecules is binary (i.e., phosphorylated or not). Importantly, the phosphorylation occupancy that we measured on multiple sites on Hec1 was similar between the soluble and insoluble (e.g., predominantly chromatin bound) populations of Hec1 in mitotic cells. Moreover, we observed partial phosphorylation occupancy on multiple sites of Hec1 in mitotic cells treated with high doses of nocodazole, conditions where all kinetochores lack any microtubule attachments and are in an equivalent state. Thus, fractional phospho-occupancy reflects equal distribution of the phosphorylated Hec1 amongst the total population of Hec1 molecules at the time of assessment as opposed to reflecting different sub-populations of Hec1 molecules where phospho-occupancy depends on the nature of the sub-population. In other words, in a case where a specific site demonstrates 25% phospho-occupancy, our data indicates that 25% of all Hec1 molecules are phosphorylated rather than specific sub-populations of Hec1 molecules where one sub-population is comprised of 25% of Hec1 molecules that are 100% phospho-occupied at that site and another sub-population that is comprised of 75% of Hec1 molecules that are 0% phospho-occupied at that site.

These findings carry numerous implications for how phosphorylation regulates kinetochore-microtubule attachments at the binding interface to both favor efficient microtubule capture by kinetochores and to support robust error correction to ensure high mitotic fidelity (Figure 7G). First, these data underscore the principle that net phosphorylation occupancy of any specific site on a protein is determined by ongoing activities of both kinases and phosphatases, and illustrates the essential role for ongoing protein phosphatase activity in regulating k-MT attachments. Specifically, some models predict that phospho-occupancy of proteins like Hec1 at the kinetochore-microtubule interface would be nearly fully saturated under conditions where kinetochores lack microtubule attachments and/or possess improperly oriented microtubule attachments ^4, 13, 34^. However, we demonstrate that in mitotic cells treated with low dose nocodazole which induces the formation of stub-spindles replete with erroneous k-MT attachments lacking tension that ongoing phosphatase activity limits the extent of Hec1 phosphorylation (Fig 6). This ongoing phosphatase activity is essential to bias kinetochores to remain attached to the stub spindles because chromosomes detach from the stub-spindles upon phosphatase inhibition. Indeed, even when kinetochores completely lack microtubule attachments induced by treatment of mitotic cells with high dose nocodazole the ongoing phosphatase activity maintains a relatively low level of Hec1 phospho-occupancy (Figure 1B, Figure 6A-D, Figure S2E). In support of this idea, it has been shown that mitotic cells in which phosphatases have been inactivated are unable to properly congress chromosomes to the metaphase plate ^10, 11^.

These observations resolve a previously described paradox ^13, 34–36^ that explicitly predicted high kinetochore phospho-occupancy in the absence of microtubule attachments. Resolving this paradox is particularly important to illuminate how conditions favor initial k-MT attachments at the prophase to prometaphase transition. At the time of nuclear envelope breakdown, kinetochores lack microtubule attachments. If kinase activity were unconstrained toward substrates on unattached kinetochores, then complete phosphorylation of sites at the prophase to prometaphase transition would substantially disfavor the formation of initial microtubule attachments. Our data highlights the importance of ongoing phosphatase activity to dampen kinetochore substrate phosphorylation in order to favor initial microtubule attachments.

We also provide the first direct evidence that Cdk1-Cyclin B1 plays an active role in promoting k-MT attachment error correction, and that Cdk1-mediated error correction is independent from the established role of Aurora kinase. Our data show that these two enzymes recognize different types of k-MT attachment errors with Cdk1 focused on conversion of sidewall to end-on attachments and Aurora focused k-MT attachments with improper tension. Consistently, sites phosphorylated by these enzymes show opposing changes in phospho-occupancy in response to the same stimulus (Figure S7E, S7F). This supports the view that different types of k-MT attachment errors rely on different mechanisms of detection and/or response to catalyze their correction. Perhaps it is surprising that these two kinases, that are responding to different types of k-MT attachment defects, phosphorylate closely located sites on the same peptide of Hec1. However, it is important to emphasize that the functional output of these kinases is not expected to be limited to changes in phosphorylation of only Hec1 at kinetochores.

This new role of Cdk1-Cyclin B1 in error correction fits within a feed-forward mechanism of mitotic control whereby Cdk1-Cyclin B1 targeting to kinetochores through Mad1 ^25, 37, 38^ regulates cell cycle timing through satisfaction of the spindle assembly checkpoint and also ensures efficient correction of erroneous k-MT attachments. Notably, the targeting of Cdk1-Cyclin B1 to kinetochores with attachment errors in early mitosis depends on Cdk1-Cyclin A2. Previously, we demonstrated that Cdk1-Cyclin A2 activity destabilizes k-MT attachment stability coordinately on kinetochores of all chromosomes to promote error correction in prometaphase ^24^ which reinforces this feed-forward mechanism involving cell cycle timing with k-MT attachment error correction. The data presented here provides further molecular insight into how a distributive system that regulates all kinetochores coordinately through Cdk1-Cyclin A2 forms direct mechanistic links to regulatory molecules that operate chromosome-autonomously (Cyclin B1 & MYPT1/Plk1) ^39^.

Finally, no site on Hec1 that we measured exceeded 50% phospho-occupancy under any condition that we tested and the cumulative phospho-occupancy among the sites measured changed by only 20% in response to the full spectrum of microtubule attachment status from unattached, to predominantly erroneous attachments, to predominantly end-on attachment. These data indicate that the correction of microtubule attachment errors responds to very small changes in phosphorylation, at least with respect to the phospho-occupancy of Hec1. As a mechanism of control, this indicates that phosphorylation displays properties of high sensitivity at the kinetochore-microtubule binding interface because small changes in phosphorylation are associated with substantial changes (i.e. detachment) in microtubule attachment status. In this context, the system appears strongly biased to maintaining inherently low phospho-occupancy which is important to both promote initial microtubule attachments and microtubule re-attachment following a detachment event related to the correction of an improperly oriented microtubule. Our data provide direct biochemical evidence for results based on amino acid substitution mutants indicating that 1-2 phosphates per Hec1 molecule are sufficient to recapitulate normal kinetochore/chromosome behavior, and that kinetochore behavior is most sensitive between 0-4 phosphates ^6^. An important distinction, however, is that we demonstrate that Hec1 molecules are phosphorylated at partial phospho-occupancy over several sites, rather than an all-or-none phospho-occupancy at 1-2 sites. Tangibly, stoichiometric assessments show that single microtubule ends are surrounded by eight Hec1 molecules at kinetochores ^40^, and our data indicates that phosphorylation of as few as three of the eight molecules of Hec1 surrounding a given microtubule (37.5% phospho-occupancy) would be sufficient to favor detachment of that microtubule from the kinetochore.

## Methods

### Cell Culture

HeLa cells (CCL-2) and RPE1 (CRL-4000) were obtained from the ATCC. All cells were grown at 37°C in a humidified environment with 5% CO_2_. The unmodified cell lines were grown in Dulbecco’s modified Eagle medium (Corning® #15-017-CM) containing 10% FCS (Hyclone #82013-586), 250μg/L Amphotericin B (VWR #82026-728), 50 U/mL penicillin and 50 μg/mL streptomycin (ThermoFisher Scientific #15140122). Doxycycline inducible Hec1 knockout HeLa cells (gift of Iain Cheeseman) were grown in media containing 10% tetracycline-free FBS (Denville Scientific #1005807), the same antibiotics as for unmodified cells, 500 μg/ml G418 (Invivogen #ant-gn-5) and 5 μg/ml puromycin (ThermoFisher Scientific #A1113803). HeLa Cyclin B1-GFP cells were grown in media containing 4 μg/ml blasticidin (Invivogen # ant-bl-05). HeLa Cyclin A2-GFP cells ^25^ were grown in media containing 9 μg/ml blasticidin (gifts of Francis Barr). Cells were verified to be free of mycoplasma by frequent staining of plated cells with DAPI. No contamination was observed.

### Inhibitors and reagents

The CENP-E inhibitor GSK-923295 (MedChemExpress LLC # HY-10299) was used at 200 nM. NOC (VWR #80058-500) was used at 100-500 ng/ml. STLC (Tocris #2191) was used at 25 μM. ProTAME (Concept Life Sciences custom synthesis) was used at 25 μM. Okadaic acid (LC Labs O-5857) was used at 200 nM. Doxycycline hyclate (Sigma-Aldrich #24390-14-5) was used at 2 μg/ml.

### Plasmids and cloning

pFETCh_Donor (EMM0021) was a gift from Eric Mendenhall & Richard M. Myers (Addgene plasmid # 63934). pSpCas9(BB)-2A-GFP (PX458) was a gift from Feng Zhang (Addgene plasmid # 48138). The Hec1-GFP Wt and 9A mutant plasmids were a gift of Jennifer DeLuca. Additional mutations were generated via site-directed mutagenesis using the following primers: S44 Wt/8A FD: 5’-

GAGAAACCAACCTTTGGAAAGTTGAGTATAAACAAACCGGCCTCTGAAAGA-3’

RV: 5’-TCTTTCAGAGGCCGGTTTGTTTATACTCAACTTTCCAAAGGTTGGTTTCTC-3’

T31A FD: 5’-AAACAAGGCCTCTATGCCCCTCAAACCAAAGAGAAACC-3’

RV: 5’-GGTTTCTCTTTGGTTTGAGGGGCATAGAGGCCTTGTTT-3’

T31D FD: 5’-AAACAAGGCCTCTATGACCCTCAAACCAAAGAGAAACC-3’

RV: 5’-GGTTTCTCTTTGGTTTGAGGGTCATAGAGGCCTTGTTT-3’

S15A FD: 5’-GGTGCTGGCCGCCTCGCCATGCAGGAGTTAAGATCC-3’

RV: 5’-GGATCTTAACTCCTGCATGGCGAGGCGGCCAGCACC-3’

S15D FD: 5’-GGTGCTGGCCGCCTCGACATGCAGGAGTTAAGATCC-3’

RV: 5’-GGATCTTAACTCCTGCATGTCGAGGCGGCCAGCACC-3’

The DNA constructs were then verified by sequencing and amplified by maxiprep (Qiagen #12162)

### Transfections

100,000 HeLa inducible Hec1 knockout cells were plated on glass coverslips in 12-well plates. 24 hours following plating, they were transfected with 1 μg plasmid DNA and 2.5 μl Lipofectamine^TM^ 2000 (ThermoFisher #11668027) in a total of 1 ml Opti-MEM^TM^ media (ThermoFisher #31985070). Four hours following transfection, the media was changed for DMEM containing doxycycline, puromycin and G418. The cells were then assayed 48-72 hours following transfection.

### Mitotic error rate

100,000 HeLa inducible Hec1 knockout cells were plated on glass coverslips in 12-well plates. For determination of the error rate in asynchronous cells, the cells were fixed 72 hours after transfection, stained with DAPI and then analyzed. For determination of the error rate from STLC washout, the cells were transfected as described above, allowed to recover for 24 hours and then synchronized by thymidine addition for 24 hours. The cells were then released into media containing STLC for 14 hours. The cells were then released from STLC by two PBS washes and then fresh medium for 2.5 hours before the cells were fixed.

### CRISPR/Cas9 genomic editing

We employed a strategy described previously ^41^ to tag Hec1 at the endogenous locus (NCBI Reference Sequence: NC_000018.10). HeLa cell genomic DNA was purified using a kit (Qiagen #69504). The homology arms were then amplified by PCR using NEB Phusion® polymerase (#M0530S) using the following primers:

Hom1 FD: 5’-TCCCCGACCTGCAGCCCAGCTGATAAATACGAAGGTGAAAACCAGG-3’ Hom1 RV: 5’-CCGGAACCTCCTCCGCTCCCTTCTTCAGAAGACTTAATTAGAG-3’ Hom2 FD: 5’-AGTTCTTCTGATTCGAACATCGAATAAAATTGTCTCAGTAAAGTG-3’ Hom2 RV: 5’-TGGAGAGGACTTTCCAAGGGATTTGGCTCACATGATGTAGG-3’

A donor plasmid was then created via Gibson assembly according to the manufacturer protocol (New England Biolabs #E2611S). Separately, a vector expressing the gRNA and cas9 was created as previously described ^42^ using the following DNA fragments:

5’-CACCGTGAGACAATTTTATTCACTA-3’

5’-AAACTAGTGAATAAAATTGTCTCAC-3’

The DNA constructs were then verified by sequencing and amplified by maxiprep. 200,000 HeLa cells were then plated on a 6-well plate. 24 hours later they were transfected with 2 μg donor plasmid, 1 μg gRNA/cas9 plasmid and 7.5 μl lipofectamine^TM^ 2000 in 2 ml opti-MEM^TM^ media. Four hours Following transfection, the media was changed for DMEM. 24 hours later, G418 was added to select for successfully edited cells. Once the cells reached colonies of 50-100 cells, they were trypsinized and plated on 96-well plates at a density of 0.5 cells per well. Single-cell clones were then amplified and tested for expression of the edited gene via IF and Western blot.

### Purification of Hec1-3XFLAG

HeLa Hec1-3XFLAG cells were expanded to 16 15 cm dishes. When they reached 70-80% confluency, Thymidine (Sigma-Aldrich #T1895-5G) was added to a concentration of 2.5 mM. After 24 hours, the cells were washed twice with PBS and released into NOC or STLC. The cells were then harvested 14 hours after release. For proTAME synchronization, the cells were released from thymidine as above, but the drug was added 8 hours later for a further 4 hours before harvesting. Mitotic cells were then harvested by washing the detached and semi-detached cells off the plate by pipetting. The cells were then washed with PBS and then divided 60%-40% for subcellular fractionation and total protein purification, respectively. Cells used for total protein purification were lysed with 2 ml of a buffer containing 1% Sodium dodecyl sulfate (SDS; Sigma-Aldrich #436143), 50 mM Tris pH 7.5, 1mM EDTA. The cells were then crudely homogenized using a 5 ml syringe and 18 g needle. The lysate was then immediately boiled for 5 minutes. After cooling, the DNA was sheared using a syringe until the viscosity was acceptably reduced in a 15 ml tube using a 5ml syringe and 18g needle. The SDS was sequestered via addition of 13 ml extraction buffer (3% Triton X-100 (Sigma-Aldrich #T8787), 50 mM Tris pH 7.5, 250 mM NaCl, 1 mM EDTA) and cooled on ice. Concurrently, subcellular fractionation was performed on the remaining cells by suspending them in 10 ml cold hypotonic buffer (10 mM Tris pH 7.5, 10 mM KCl, 1 mM MgCl_2_, 10 mM glycerol 2-phosphate (Sigma-Aldrich #G9422), 1 mM EDTA) on ice, followed by douncing every 5 minutes. After 20 minutes, the mixture was aliquoted into 1.5 ml tubes and centrifuged at 4°C at maximum speed for 20 minutes. The supernatant containing soluble protein was then removed, SDS added to a concentration of 0.2% and boiled for 5 minutes. After cooling to room temperature, 5 ml of 3X extraction buffer was added to the soluble fraction, which was set aside on ice. The chromatin containing pellets were crudely homogenized via syringe in 2 ml SDS lysis buffer and boiled for 5 minutes. The DNA was then sheared via syringe in a 15 ml tube. The SDS was then sequestered by adding 13 ml extraction buffer and cooled on ice. All 3 fractions were then centrifuged at maximum speed in a benchtop centrifuged to remove any remaining cell debris from the lysates. 30 μl of packed M2 anti-FLAG affinity gel (Sigma-Aldrich #A2220-5ML) was then added to the lysates, which were then incubated overnight on a rotating platform at 4°C. The affinity gel was then transferred to a 1.5 ml tube and washed twice with extraction buffer. The volume of extraction buffer was then reduced to 30 μl, and 70 μl water and 1mM MnCl_2_ added together with 1 μl DNaseI (NEB #M0303S) for 10 minutes at room temperature in order to degrade any DNA that precipitated overnight. Any separable debris was aspirated using a 30g needle. The affinity gel was then washed an additional 3 times with cold extraction buffer, incubating for 10 minutes between each wash. The gel was then dried using a 30 g needle connected to a vacuum line. To elute the purified protein, the affinity gel was suspended in elution buffer (2% SDS, 50 mM Tris pH 7.5, 50 mM NaCl, 20% glycerol, 2 mM dithiothreitol (Sigma-Aldrich #D0632-5G; fresh) and incubated at 65°C for 20 minutes. The samples were then cooled to room temperature.

Iodoacetamide (Sigma-Aldrich #I6125-5G) was added to a concentration of 6mM, vortexed and the samples incubated in the dark for 1 hour. Then, an additional 2mM dithiothreitol was added and incubated for 15 minutes. The samples were then run on 8% SDS-PAGE gel and stained with Coomassie blue. The gels were imaged with a Gel Doc^TM^ Ez (Bio-Rad #1708270).

### SDS-PAGE and Western blot

Gels and blots were performed as described previously ^43^.

### Mass spectrometry

Gel bands were excised, destained to clarity, digested using trypsin (Promega # V5113) and the resulting peptides were extracted as described previously ^44^. Each sample was desalted by STAGE tip ^45^ and then dried by vacuum centrifugation. Samples were resuspended in 220μl of buffer (100mM TEAB (Sigma-Aldrich #17902), 10mM MgCl_2_, and 100mM NaCl) by vortexing and split into pairs of 100μl each. One of each pair was treated with 0.5 μl of calf intestinal phosphatase (CIP) (New England Biolabs #M0290) before the pair were incubated at 37°C for 1 hour. Since the Aurora kinases phosphorylate proteins adjacent to lysine and arginine residues, which also represent trypsin cleavage sites, both CIP-treated and untreated samples were then re-digested with an additional 0.1μg trypsin overnight at 37°C to prevent missed cleavages due to protein phosphorylation. The CIP-treated and untreated samples were then labeled by heavy and light reductive dimethyl labeling respectively, as previously described ^46, 47^ (Sigma-Aldrich #252549, #156159, #190090, Cambridge Isotope Laboratories #CDLM-4599-1). After labeling, the heavy and light pair were mixed, desalted together by STAGE tip, and dried by vacuum centrifugation. Dried samples were resuspended in 5% Methanol (Sigma-Aldrich #34860) / 1.5% Formic Acid (Honeywell# 64-18-6) and analyzed by online microcapillary liquid chromatography-tandem mass spectrometry using either a ThermoFisher Scientific QE +, Orbitrap Fusion, or an Orbitrap Lumos nLC-MS/MS platform as previously described ^48^. The mass spectra were searched using the COMET search algorithm ^49^ to generate peptide spectral matches as previously described for dimethyl labeled samples ^47^. Peptide spectral matches were filtered to a ∼1% FDR using the target-decoy strategy ^50^ and reported. Peptide masses and retention times were then used to manually quantify the extracted ion chromatograms of unphosphorylated heavy and light labeled pairs in all runs using Xcalibur Qual Browser software (ThermoFisher). Phosphorylation occupancy was calculated for candidate sites using these areas as described previously ^51^.

### RNA interference

Gene knockdown was performed as described previously ^43^ with the following siRNAs purchased from Sigma-Aldrich:

Cyclin B1: 5’-GACCAUGUACAUGACUGUCUC[dT][dT]-3’

Cyclin A2: 5’-UAUACCCUGGAAAGUCU[dT][dT]-3’

BubR1: 5’-AAAGAUCCUGGCUAACUGUUC[dT][dT]-3’

B56 Alpha: 5’-UGAAUGAACUGGUUGAGUA[dT][dT]-3’

B56 Beta: 5’-GAACAAUGAGUAUAUCCUA[dT][dT]-3’

B56 Gamma: 5’-GGAAGAUGAACCAACGUUA[dT][dT]-3’

B56 Delta: 5’-UGACUGAGCCGGUAAUUGU[dT][dT]-3’

B56 Epsilon: 5’-GCACAGCUGGCAUAUUGUA[dT][dT]-3’

NSC 5’-AAUUCUCCGAACGUGUCACGU[dT][dT]-3’

### Immunofluorescence

Following treatments, cells were fixed in methanol at −20°C except for experiments using GFP constructs, cold stable MT assays and those involving Cyclins A2 or B1, which were fixed with 4% paraformaldehyde in PBS for 20 minutes at room temperature. Cells were pre-extracted in PHEM buffer (60 mM PIPES, 25 mM HEPES, pH 6.9, 10 mM EGTA, and 4 mM MgSO_4_, 1% Triton X-100, 10 mM glycerol 2-phosphate) for 10 minutes at 4°C. The remaining steps were performed at room temperature. Experiments involving Cyclin A2 or Cyclin B1 were not pre-extracted and instead were permeabilized post-fixation with PHEM buffer for 10 minutes. The cells were then re-hydrated in PBS for 10 minutes before blocking with TrueBlack® IF blocking buffer (Biotium #23012B) for 10 minutes. Primary antibodies were then diluted in Trueblack® and used at the indicated dilution. Secondary antibodies were diluted in 10% FCS, 0.1% Triton X-100 in PBS. After incubation with antibodies, the cells were washed three times with PBS. Finally, the cells were counterstained with DAPI and mounted on slides using ProLong^TM^ Gold antifade reagent (ThermoFisher Scientific #P36934).

### Microscopy

Images were acquired with a Nikon Eclipse Ti microscope equipped with a cooled charge-coupled device Clara camera (Andor Technology) controlled by Nikon NIS-Elements software version 4.30.02. Images were acquired in 0.15-0.4 μm sections using a plan apo 1.4 numerical aperture 100X (kinetochore analysis) or 60X (lagging chromosome analysis) objective using 1X1 binning. Samples were illuminated using an X-cite light source (Excelitas Technologies Corp). All image analysis, adjustment and cropping was performed using Fiji software ^52^. Image deconvolution was performed using Nikon Elements batch deconvolution software version 5.21.00 on automatic mode. All kinetochore intensity analysis was performed on the raw images, except for Figures 3a-b, for which deconvolution was performed prior to analysis. Line scans were performed on the deconvoluted images in Fiji and the data was then exported to Graphpad Prism. All images selected for presentation are deconvolved maximum-intensity Z-stack projections. Where shown, insets are from single optical Z-slices. Images were selected to represent the mean quantified data. Quantification of kinetochore staining intensity was performed by first determining the brightest plane for which a given kinetochore appears. Then, an outline was drawn using the ellipse tool. The integrated intensity was determined by multiplying the average intensity by the area of the kinetochore. Background levels of staining were determined by saturating the brightness and contrast settings to find the darkest spot within the chromatin containing area and were subtracted from the intensity of the kinetochore. To quantify levels of cold-stable microtubules, images were first deconvolved. Then, A sum-intensity projection was created in Fiji. Viewing the ACA channel, a rectangle was drawn that was 2 μm wide in the pole-pole axis, and as long as the furthest centromeres in the metaphase plate. Then, switching to the tubulin channel, the average tubulin intensity was measured in the rectangle. From the average intensity value, the minimum intensity value was used as the background level and subtracted. The Background-subtracted intensity values were then averaged across all of the cells that were analyzed for each condition and experimental repeat.

### Antibodies

The following antibodies were used for immunofluorescence (IF) and/or immunoblotting (IB): ACA (Geisel School of Medicine; IF at 1:2000), Hec1 (Santa Cruz C-11; IF,IB at 1:1000), PP2A B56 Alpha (MyBioSource.com MBS8524809; IB at 1:500), PP2A B56 Beta (Santa Cruz E-6; IB at 1:500), PP2A B56 Gamma (Santa Cruz A-11; IB at 1:500), PP2A B56 Delta (Santa Cruz H-11; IB at 1:500), PP2A B56 Epsilon (Aviva Systems Biology RP56694_P050; IB at 1:500), Nuf2 (Santa Cruz E-6; IB at 1:1000), phospho-Hec1 Ser44 (gift of Jennifer DeLuca; IF at 1:500), GAPDH (Santa Cruz G-9; IB at 1:1000), phospho-Histone H3 Ser10 (Cell Signaling Technologies #9701; IB at 1:2000), FLAG M2 (Sigma-Aldrich; IF and IB at 1:1000), Cyclin A2 (Genetex GT2547; IF at 1:1000), Cyclin B1 (Santa Cruz GNS1; IF at 1:500), α-Tubulin DM1α (Sigma-Aldrich; IF at 1:4000). The phospho-Hec1 Thr31 antibody was generated by Pacific Immunology where two rabbits were immunized with the peptide corresponding to amino acids 25-37: Cys-NKQGLY-pT-PQTKEK, followed by affinity purification against the non-phospho and phospho peptides (IF at 1:100,000). Secondary antibodies used were highly cross-adsorbed Alexa Fluor® 488, 594 and 647 (Invitrogen; IF at 1:1000) and horseradish peroxidase (Bio-Rad; IB at 1:1000-10,000), Mouse TrueBlot® ULTRA: Anti-Mouse Ig HR (Rockland 18-8817-33; IB at 1:1000).

### Statistical analysis

All statistical tests were performed using Graphpad Prism version 8.4.2. Except where indicated, all experiments were performed as 3 independent biological repeats. In most IF experiments, 20 cells per repeat were analyzed, and 25 kinetochores for each cell measured. Experiments with other numbers are noted in the figure legends. For anaphase error rates, 120 anaphases per condition were assessed for each of 3 independent experiments. Statistical significance was calculated between indicated conditions using two-tailed *t*-tests.

### Data availability

The cell biology datasets generated during and/or analysed during the current study are available from the corresponding author on reasonable request. The MS dataset presented in Figure 1 and Extended data 2 will be uploaded to an appropriate public data repository upon acceptance of the article.

## Acknowledgements

We thank Jennifer DeLuca, Francis Barr and Iain Cheeseman for providing reagents. We also thank the members of the Compton laboratory for helpful discussions and feedback. We thank Ann Lavanway, Zdenek Svindrych and the the Dartmouth College Life Sciences Light Microscopy Facility for assistance with microscopy and image deconvolution. TJK was supported by fellowships from the Fonds de la Recherche en Santé du Québec #35556 and the Canadian Institute of Health Research # 395782. This work was also supported by grants from the National Institutes of Health # P20-GM113132 to the BioMT facility of Dartmouth College, #GM122846 to SAG, #1R01HD101436 to KMG and #GM051542 to DAC.

## Author Contributions

TJK conceived the project with together with DAC, prepared all biological samples, performed all cell biology experiments and cloning, analyzed the data, prepared the figures, wrote the original manuscript draft and participated in manuscript editing. RH refined the MS protocol, performed the MS analysis and performed the initial MS quantifications. KMG provided guidance and advice and participated in manuscript editing. SAG supervised RH, assisted with the MS development, provided reagents and advice and participated in manuscript editing. DAC conceived the project with TJK, reviewed and edited the figures and manuscript and was responsible for overall supervision of the project.

## Materials & Correspondence

Any requests should be addressed to DAC

---

**Figure S1 related to** Figure 1**: Addition of a 3XFLAG tag to Hec1 does not perturb mitosis**

A. Schematic showing the addition of a 3XFLAG-P2A-Neo casette to the 3’ of the endogenous Hec1 gene. The blue highlight indicates the original stop codon.
B. Immunofluorescence images of HeLa Hec1-3XFLAG and parental HeLa cells. The cells were fixed and stained for FLAG, ACA and DAPI. The FLAG and ACA channels were adjusted evenly for brightness and contrast for presentation. The DAPI channel was adjusted independently. Representative images from 2 independent experiments are shown. The scale bars are 10 μm.
C. Immunofluorescence images of HeLa Hec1-3XFLAG and parental HeLa cells. The cells were fixed and stained for Hec1 and DAPI. The Hec1 and DAPI channels were adjusted evenly for brightness and contrast for presentation. Representative images from 3 independent experiments are shown. The scale bars are 10 μm.
D. Quantification of the percentage of lagging chromosomes in cells undergoing anaphase from an asynchronous population of HeLa Hec1-3XFLAG and parental HeLa cells. 100 cells per cell line were scored for the presence of lagging chromosomes for each of 3 independent experiments. Error bars indicate the mean +/- SEM.
E. Quantification of the time spent in mitosis from the onset of cell rounding to anaphase from an asynchronous population of HeLa Hec1-3XFLAG and parental HeLa cells. 50 cells for each population were quantified from a single experiment. Error bars indicate the mean +/- SEM.
F. Western blots showing an anti-FLAG immunoprecipitation from HeLa Hec1-3XFLAG and parental HeLa cells. The cells were synchronized by thymidine-NOC and then lysed in NETN buffer. Hec1-3XFLAG was then immunoprecipitated, and the purified proteins then separated by SDS-PAGE, transferred to nitrocellulose membrane and finally blotted for Nuf2 and FLAG. A single experiment was performed.

**Figure S2 related to** Figure 1**: Hec1 is purified from cells treated to generate different types of k-MT attachment; Both soluble and chromatin localized Hec1 is only partially phosphorylated**

A. Immunofluorescence images of HeLa Hec1-3XFLAG cells synchronized by thymidine and then NOC, STLC or proTAME. The cells were fixed and stained for ACA, tubulin and DAPI. The tubulin and ACA channels were adjusted evenly for brightness and contrast for presentation. DAPI channel was adjusted independently for presentation. The scale bars are 10 μm. Representative images from a single experiment are shown.
B. Quantification of inter-centromere distances of cells prepared as in panel (A). 100 pairs of centromeres were measured across 10 cells from a single experiment.
C. Western blots showing an anti-FLAG immunoprecipitation from HeLa Hec1-3XFLAG and parental HeLa cells. The Hela-3XFLAG cells were synchronized by thymidine-NOC and then prepared as a total cell lysate or fractionated into soluble and chromatin fractions. Control parental HeLa cells remained asynchronous and were prepared as a total cell lysate. Hec1-3XFLAG was then immunoprecipitated and the purified proteins were then separated by SDS-PAGE, transferred to nitrocellulose membrane and finally blotted as indicated. The upper band of the Hec1 (endogenous epitope) input blot represents Hec1-3XFLAG, while the lower band represents unmodified Hec1. * indicates a non-specific band. The ponceau stained panel was adjusted for brightness and contrast. A single experiment was performed.
D. Cartoon illustration depicting the calculation of absolute phosphorylation occupancy. The ratio of the unphosphorylated peptide (colored magenta) between the heavy and light channels is used to calculate phosphorylation occupancy.
E. Histograms showing the absolute phosphorylation occupancy of various Hec1 peptides purified from cells fractionated into soluble and chromatin fractions as determined by MS. Individual and average values from 3 independent experiments are shown. The subcellular fractions are derived from the same cells as used for the total protein analysis presented in figure 1. Error bars indicate mean +/- SEM. * denotes p < 0.05, ** denotes p < 0.01.
F. MS peaks for the Hec1 peptide LSINKPTSER purified from the chromatin fraction of cells synchronized by thymidine then 100 ng/ml NOC. A representative trace from one of the three replicate experiments presented in Extended data 2e is shown. The magenta trace indicates the CIP treated, heavy labelled peptides and the blue trace indicates the sham treated light labelled peptides. The absence of peaks in the magenta trace indicates complete removal of phosphates.

**Figure S3 related to** Figure 2**: The antibody raised against pT31 binds only to pT31; Hec1 is phosphorylated on T31 in prometaphase RPE1 cells; pT31 is maximally phosphorylated in cells arrested in STLC when analyzed by IF; Hec1 pT31 is antagonized by protein phosphatases**

A. Immunofluorescence images of HeLa doxycycline inducible Hec1 knockout cells in early prometaphase transfected with the indicated Hec1-GFP constructs. 72 hours post transfection the cells were fixed and stained for pT31 Hec1 and DAPI. The GFP and pT31 channels were adjusted evenly for brightness and contrast. The DAPI channel was adjusted independently. The scale bars are 10 μm. Representative images of 2 independent experiments are shown.
B. Immunofluorescence images of RPE1 cells in various stages of mitosis taken from an asynchronous population. 48 hours after plating the cells were fixed and stained for Hec1, pT31 Hec1 and DAPI. The Hec1 and pT31 Hec1 channels were adjusted evenly for brightness and contrast. The DAPI channel was adjusted independently. The scale bars are 10 μm. Representative images of 2 independent experiments are shown.
C. Quantification of the relative kinetochore intensities from the conditions in panel (B). The condition with the lowest level of pT31 was set to 1 and the other conditions shown as fold-changes. 25 kinetochores were quantified from each of 20 cells for each of 2 independent repeats. Error bars indicate the mean +/- SEM. **** denotes p < 0.0001.
D. Immunofluorescence images of HeLa cells synchronized by thymidine and then NOC, STLC or proTAME. The cells were then fixed and stained for Hec1, pT31 Hec1 and DAPI. The Hec1 and pT31 Hec1 channels were adjusted evenly for brightness and contrast. The DAPI channel was adjusted independently. The scale bars are 10 μm. Representative images of 3 independent experiments are shown.
E. Quantification of the relative kinetochore intensities from the conditions in panel (B). The condition with the lowest level of pT31 was set to 1 and the other conditions shown as fold-changes. 25 kinetochores were quantified from each of 20 cells for each of 2 independent repeats. Error bars indicate the mean +/- SEM. **** denotes p < 0.0001.

**Figure S4 related to** Figure 4**: Cyclin B1 only localizes strongly to kinetochores that have improper k-MT attachments**

A. Immunofluorescence images from HeLa Cyclin B1-GFP cells synchronized with thymidine and then NOC, STLC or proTAME. The cells were then fixed and stained for Hec1, pT31 Hec1 and DAPI. The Hec1 and Hec1 pT31 channels were adjusted evenly for brightness and contrast. The GFP and DAPI channels were adjusted independently. The scale bars for the main images are 10 μm and 1 μm for the insets. Representative images of 2 independent experiments are shown.
B. Line scans through the indicated pairs of kinetochores from the cells shown in panel (A).

**Figure S5 related to** Figure 5**: Hec1 is phosphorylated at T31 on unaligned chromosomes**

A. Immunofluorescence images of HeLa cells in late prometaphase from an asynchronous population treated with GSK-923295 for 2 hours and then fixed and stained for Hec1, pT31 Hec1 and DAPI. The images were adjusted for brightness and contrast for presentation. The top insets show aligned kinetochores, and the lower insets show unaligned kinetochores. The scale bars for the main images are 10 μm and 1 μm for the insets. Representative images of 3 independent experiments are shown.
B. Quantification of the relative kinetochore intensities from the conditions in (A). The levels of the aligned kinetochores were set to 1 and the intensity of the unaligned kinetochores shown as a fold-change. 1509 (aligned) and 1355 (unaligned) total kinetochores were quantified from 20 cells for each of 3 independent repeats. Error bars indicate the mean +/- SEM. **** denotes p < 0.0001.
C. Immunofluorescence images of HeLa doxycycline inducible Hec1 knockout cells transfected with the Hec1-8A construct. The cells are in late prometaphase taken from an asynchronous population. 72 hours post transfection the cells were fixed and stained for pT31 Hec1 and DAPI. The GFP and pT31 channels were adjusted evenly for brightness and contrast. The DAPI channel was adjusted independently. The top inset shows aligned kinetochores and the bottom inset shows unaligned kinetochores. The scale bars are 10 μm. Representative images of 3 independent experiments are shown. The cells presented in this experiment are the same experiment presented in Figure S7A but re-analyzed to compare the levels of pT31 in unaligned vs aligned kinetochores.
D. Quantification of the relative kinetochore intensities from the conditions in (C). The levels of the aligned kinetochores were set to 1 and the intensity of the unaligned kinetochores shown as a fold-change. 957 (aligned) and 625 (unaligned) total kinetochores were quantified from at least 10 cells for each of 3 independent repeats. Error bars indicate the mean +/- SEM. **** denotes p < 0.0001. The cells analyzed in this experiment are the same cells presented in Figure S7A but re-analyzed to compare the levels of pT31 in unaligned vs aligned kinetochores.
E. HeLa Cyclin B1-GFP cells were treated as in panel (A), but then stained for ACA, tubulin and DAPI. The images were adjusted for brightness and contrast for presentation. The scale bars for the main images are 10 μm and 1 μm for the insets. Representative images of 2 independent experiments are shown.
F. Line scans through the kinetochores shown in the insets.

**Figure S6 related to** Figure 7**: Depletion of the BubR1 pool of PP2A-B56 phosphatase increases Hec1 phosphorylation and decreases k-MT stability**

A. Immunofluorescence images from an asynchronous population of HeLa cells transfected with control or siRNA against BubR1 and then stained for Hec1 pS44, Hec1 and DAPI. The Hec1 and Hec1 pS44 channels were adjusted evenly for brightness and contrast. The DAPI channel was adjusted independently. The scale bars for main images are 10 μm and 1 μm for insets. Representative images of 3 independent experiments are shown.
B. Quantification of the relative kinetochore intensities from the conditions in panel (A). The condition with the lowest level of pS44 was set to 1 and the other conditions shown as fold-changes. 25 kinetochores were quantified from each of 20 cells for each of 3 independent repeats. Error bars indicate the mean +/- SEM. **** denotes p < 0.0001.
C. Immunofluorescence images of cells prepared as in panel (A) except stained for pT31 instead of pS44. Representative images of 3 independent experiments are shown.
D. Quantification of the relative kinetochore intensities from the conditions in panel (B). The condition with the lowest level of pT31 was set to 1 and the other conditions shown as fold-changes. 25 kinetochores were quantified from each of 20 cells for each of 3 independent repeats. Error bars indicate the mean +/- SEM. **** denotes p < 0.0001.
E. Immunofluorescence images from an asynchronous population of HeLa cells transfected with control or siRNA against BubR1. The media was then changed for media at 4°C, and the cells placed at 4°C for 15 minutes prior to fixation and staining with antibodies against ACA and α Tubulin. Representative images of 3 independent experiments are shown.
F. Quantification of the relative intensity of α Tubulin. The levels of the control condition were set to 1 and the other condition shown as a fold-change. At least 50 cells per condition from each of 3 independent experiments were quantified. Error bars indicate the mean +/- SEM. **** denotes p < 0.0001.
G. HeLa Cyclin B1-GFP cells were transfected with control or a pool of siRNA against the 5 B56 isoforms. The cells were then fixed and stained for ACA, tubulin and DAPI. The channels were adjusted independently for presentation. Representative images of 3 independent experiments are shown.
H. Quantification of the percentage of cells displaying Cyclin B1-GFP localization at kinetochores from panel (G). At least 20 cells were imaged per experimental condition for each of 3 independent experiments. The cells with B56 knockdown were identified as having either moderate or severe alignment defects. All cells were scored for having Cyclin B1 at kinetochores or not. Error bars indicate the mean +/- SEM. *** denotes p < 0.001.
I. HeLa Cyclin B1-GFP cells were transfected with control or siRNA against BubR1. The cells were then fixed and stained for ACA, tubulin and DAPI. The channels were adjusted independently for presentation. Representative images of 3 independent experiments are shown.
J. Quantification of the percentage of cells displaying Cyclin B1-GFP localization at kinetochores from panel (I). At least 20 cells were imaged per experimental condition for each of 3 independent experiments. All cells were scored for having Cyclin B1 at kinetochores or not. Error bars indicate the mean +/- SEM. ** denotes p < 0.01.
K. HeLa cells transfected with control or siRNA against BubR1. They were then harvested, lysed and run on SDS-PAGE and transferred onto membrane. The membranes were then blotted as indicated.
L. HeLa cells transfected with control or a pool of siRNA against B56 α-ε. They were then harvested, lysed and run on SDS-PAGE and transferred onto membranes. The membranes were then blotted as indicated.

**Figure S7 related to** Figure 7**: pT31 and pS44 are differentially regulated Schematic of Hec1 mutants; Inter-kinetochore distance is increased in cells expressing an 8A mutant and decreased in cells expressing an 8D mutant compared to Wt Hec1**

A. Cartoon illustration of the Hec1-GFP constructs used in Figures S5 and S7 showing the mutations made and the remaining potential phosphorylation sites.
B. Quantification of inter-kinetochore distances of cells shown in Figure S5 and S7. 660, 650, 640, 660,660 and 670 total pairs of kinetochores for the Wt, 8A, 8D (early prometaphase) and Wt, 8A, 8D (late prometaphase), respectively were measured across 20 cells in each of 3 independent experiments. Error bars indicate the mean +/- SEM. * denotes p < 0.05, *** denotes p < 0.001, **** denotes p < 0.0001.
C. Immunofluorescence images of HeLa doxycycline inducible Hec1 knockout cells transfected with the indicated Hec1-GFP constructs. 72 hours post transfection the cells were fixed and stained for pS44 Hec1 and DAPI. The GFP and pS44 Hec1 channels were adjusted evenly for brightness and contrast for presentation. The DAPI channel was adjusted independently. Representative images from 3 independent experiments are shown. The scale bars are 10 μm.
D. As for (A) except that the cells were stained for pT31 instead of pS44.
E. Quantification of the relative kinetochore intensities from the conditions in (C). The condition with the lowest level of pS44 was set to 1 and the other conditions shown as fold-changes. 25 kinetochores were quantified from each of 20 cells for each of 3 independent repeats. Error bars indicate the mean +/- SEM. ** denotes p < 0.01, **** denotes p < 0.0001.
F. As for (E) but for pT31 instead of pS44.

